# The shrub *Ephedra californica* facilitates arthropod communities along a regional desert climatic gradient

**DOI:** 10.1101/2020.08.25.266452

**Authors:** Jenna Braun, Michael Westphal, Christopher J. Lortie

## Abstract

Arthropods underpin arid community dynamics and provide many key ecosystem services. In arid ecosystems, the key habitat components that influence arthropod community structure are relatively understudied. *Ephedra californica* is a common and widespread shrub with established positive effects on plant and vertebrate animal communities within the drylands of Southern California. The capacity for these positive effects to further support arthropod communities has not been examined. We tested the hypothesis that the physical structure and cover vegetation enhances key measures of arthropod community assembly at nine Californian desert sites that comprise an extensive regional aridity gradient. We contrasted the effects of shrub canopies with ground-covering vegetation on structuring ground-active arthropod communities by surveying ground-active arthropods with pitfall traps and collecting vegetation on the soil surface in the form of residual dry matter (RDM). We collected a total of 5868 individual arthropod specimens for a total of 184 morphospecies. Arthropod abundance and morphospecies richness and RDM biomass and cover were significantly greater beneath the canopy of *E. californica* throughout the region. Total biomass of RDM did not significantly influence arthropod communities, but cover of RDM on the soil surface negatively influenced arthropod abundance. Neither climatic aridity nor downscaled evaporative stress estimates were significant mediators of the arthropod-vegetation association patterns. Vegetation thus likely has direct and indirect physical effects on arthropod communities. These canopy versus soil surface vegetation differences will refine sampling of fine-scale patterns of arthropod diversity in drylands. Regional land managers can support arthropod diversity by maintaining populations of foundation shrub species such as *E. californica*.

## Introduction

Arthropods are the dominant component of biodiversity in arid ecosystems. Virtually every process in dryland systems is influenced by the mediation that arthropods provide through resource flows and community structure (Whitford 2000). This includes numerous key ecosystem services and trophic connections. Ground-active arthropods strongly influence ecosystem nutrient cycling through their roles as dominant detritivores (Prather et al. 2013). Ground-active beetles and spiders are important biological controls for pest species as key consumers through predation on many other species of arthropods (Suenaga and Hamamura 2001, Rosas-Ramos et al. 2018). Seed dispersal services by granivorous ants can also be critical in arid and semiarid ecosystems because they influence the structure of vegetation communities (Hulme 1998, Leal et al. 2007). Arthropods comprise a critical component of the diet of many vertebrate species under active management in California. For instance, ground-active arthropods including Coleoptera and Orthoptera comprise most of the diet of the endangered lizard species *Gambelia sila* in the San Joaquin Desert of California (Germano et al. 2007). Protein content in insects is also critical to sage-grouse development in early life (Johnson and Boyce 1990). Consequently, arthropod assemblages are both a potential indicator of general ecosystem health (Kremen et al. 1993) and a fundamental taxa that warrants description and examination of drivers relevant to community dynamics. Insect community dynamics are thus highly relevant to enable support of biodiversity regionally and to manage vertebrate species at risk that rely on this taxa.

In arid and semi-arid environments, shrubs are foundation species because they commonly support biodiversity. Microclimate amelioration by shrubs is a well-established mechanism of facilitation that affects survival, growth, and reproduction of annual plants (Flores and Jurado 2003, Filazzola and Lortie 2014, McIntire and Fajardo 2014). At small scales, annual plant productivity is also often associated with higher abundances and diversity of arthropods (Siemann 1998). Vegetation provides physical structure as well as the nutritional base for grassland arthropod food webs (Joern and Laws 2013). Generalist predators such as carabid beetles benefit from increased density of herbivorous or detritivorous prey within vegetation and mulch (Mathews et al. 2004, Birkhofer et al. 2008). A growing body of evidence further demonstrates the importance of shrub canopy facilitation for arthropod communities in arid ecosystems (Groner and Ayal 2001, Sanchez and Parmenter 2002, Liu et al. 2014, Liu et al. 2016, Ruttan et al. 2016, Braun and Lortie 2020). Therefore, there is the capacity for direct positive effects on arthropods by shrubs, and there is also capacity for indirect effects mediated through benefits to the annual plant understory. However, these indirect canopy and facilitation effects of shrubs on other species through vegetation is rarely examined within these studies. In arid environments the growing season for annual plants is very short. Residual dry matter (RDM) is the standing annual plant biomass, and it is a standard method used by land managers to quantify grazing pressure (Bartolome et al. 2002). RDM is a persistent component of the system because decomposition is relatively slow with limited precipitation (Robinson et al. 1995). In California, the activity period of ground arthropods extends beyond the end of the annual growing season (e.g.VanTassel et al. 2014); however, the influence of RDM on arthropod communities has not been examined. RDM can thus in theory support arthropod communities by providing additional habitat structure or refuges. Alternatively, RDM can physically interfere with arthropods as it does for some small vertebrates by limiting movement (Germano et al. 2012, Filazzola et al. 2017). Understanding the relative importance of shrubs and RDM on the structure of arthropod communities will inform the ecological and habitat needs of ground-active arthropod communities.

Direct and indirect effects of shrubs on plants can change depending on precipitation, relative stress, and spatially at many scales on gradients. The stress gradient hypothesis (SGH) proposes that the balance of plant-plant interactions shifts to facilitative or positive under increasing stress due to habitat amelioration (Bertness and Callway 1994), and these increases in frequency can expand the realized niche for some of the beneficiary species (He and Bertness 2014). Shrub canopy facilitation can increase in frequency along a stress gradient by providing relatively more stable micro-climates (Pugnaire et al. 2011). Extensions of the SGH to other trophic levels focus on interactions between animals using the same set of resources, such as aquatic detritivores (Fugère et al. 2012) and herbivores (Barrio et al. 2013, Dangles et al. 2013). It is not known if the facilitative effects of shrubs on arthropod communities increase in relative ecological importance or in frequency as relative environmental stress increases. Climate models predict an increase in the intensity and frequency of droughts and water stress in drylands in California, and globally (MacDonald 2007, Solomon et al. 2007). Understanding how these interactions change under stress can help predict the outcomes of global change on the structure and biodiversity of affected communities (Callaway 2007).

*Ephedra californica* is a locally abundant shrub now restricted to highly fragmented populations regionally within California deserts that has demonstrated positive effects on plant communities (Filazzola et al. 2017, Lortie et al. 2018). The capacity for these positive effects to extend to arthropod communities has not been tested. Here, we measured the association patterns of arthropod communities with shrubs and measured RDM on a regional aridity gradient within the central drylands of California. We chose early summer because it is a period of high activity for several species of endangered vertebrate animals that are under management within these ecosystems that depend on arthropods for a major component of their diets (Germano et al. 1994, Germano et al. 2007). We examined the hypothesis that the shrub species *Ephedra californica* and RDM jointly influence the structure of ground-active arthropod communities by testing the following predictions.

1. Shrubs facilitate the annual plant community directly through relative increases in biomass and cover of RDM.
2. Shrubs consistently facilitate biodiversity by increasing arthropod abundance and morphospecies richness relative to paired non-canopied microsites.
3. RDM influences the arthropod community and interacts with the canopy effects of shrubs.
4. Increasing aridity and environmental stress on a regional gradient increases the positive effects of vegetation on key measures of the arthropod community.

These predictions provide an assessment of the relative importance of canopy and ground vegetation in supporting arthropod communities in drylands and thus biodiversity of taxa critical for supporting ecosystem functioning. There is significant pressure in these regions through extended droughts (Kogan and Guo 2015) and land use changes; however, the native shrub species *Ephedra californica* is still widely distributed in the region (Calflora 2020). Another objective was to evaluate the consistency of the composition of arthropod assemblages associated with *E. californica.* If *E. californica’*s indicator species are consistent throughout the extent of the study area, then associated arthropod communities are likely specialized to the shrub *E. californica.* However, if the indicator species are different, that will indicate that *E. californica* has more generalized, positive effects that regionally enhance biodiversity in addition to its finer scale effects.

## Methods

### Site description

We sampled nine sites across semi-arid and arid desert regions forming an aridity gradient in Southern California, U.S.A between the dates of June 23^rd^ and July 8^th^, 2019 (Table 1, Appendix A1). The shrub species *Ephedra californica* (Ephedraceae) is the dominant perennial species with the semi-arid extents of the site gradient at Panoche Hills and Cuyama Valley within the San Joaquin Desert, and it is codominant with *Larrea tridentata* (Zygophyllaceae) at arid sites in the Mojave Desert. All sites are administered by the Bureau of Land Management. Soil composition at Panoche Hills is well-drained and varies from sandy loam to gravely loam dependent on slope (Arroues 2006). Mojave sites are sandy and typical for *Larrea tridentata* (Lei 1998). Site-level mean annual temperature, mean annual precipitation and the maximum temperature of the warmest month were obtained from WorldClim (Fick and Hijmans 2017). DeMartonnes aridity values were calculated using the following formula: 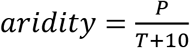 where P = annual precipitation and T = mean annual temperature. The evaporative stress indicator (ESI) was retrieved from the ECOSTRESS sensor from the daytime dates most closely corresponding to the trapping (Appendix A2). We used the point value corresponding with the coordinates for the site (Table 1), the data is provided at 70 m spatial resolution. ESI is evapotranspiration/potential evapotranspiration and is a local short-term measure of water deficit for plants (Meerdink et al. 2019), whereas aridity is a measure of overall climate.

**Table 1:** List of study sites surveyed for this project. Mean annual temperature (MAT) and mean annual precipitation (MAP) extracted from WorldClim and used to calculate DeMartonnes Aridity Index. Evaporative stress index (ESI) from EcoSTRESS.

| Regional<br>Gradient<br>Classification | Site | Desert | Latitude | Longitude | Elevation<br>(m) | MAT<br>(°C) | MAP<br>(mm) | Aridity | ESI |
| --- | --- | --- | --- | --- | --- | --- | --- | --- | --- |
| Semi-arid | 1 | San Joaquin | 34.84873 | -119.483 | 848 | 14 | 547 | 22.792 | 0.114 |
| Semi-arid | 2 | San Joaquin | 34.85362 | -119.486 | 837 | 14.1 | 542 | 22.490 | 0.085 |
| Semi-arid | 3 | San Joaquin | 34.93824 | -119.481 | 827 | 14.2 | 513 | 21.198 | 0.087 |
| Semi-arid | 4 | San Joaquin | 36.70001 | -120.801 | 656 | 14.4 | 381 | 15.615 | 0.226 |
| Semi-arid | 5 | San Joaquin | 36.70654 | -120.812 | 611 | 14.4 | 377 | 15.451 | 0.222 |
| Semi-arid | 6 | San Joaquin | 36.70554 | -120.812 | 596 | 14.4 | 377 | 15.451 | 0.236 |
| Arid | 7 | Mojave Desert | 34.6982 | -115.684 | 784 | 19.7 | 135 | 4.545 | 0.034 |
| Arid | 8 | Mojave Desert | 34.20568 | -115.72 | 545 | 20.9 | 109 | 3.528 | 0.067 |
| Arid | 9 | Mojave Desert | 35.09405 | -116.835 | 496 | 19.3 | 79 | 2.696 | 0.023 |

### Vegetation sampling

At each study site, we selected 30 pairs of shrub and open microsites. Shrub microsites were surveyed on the north side of the shrub within the canopy dripline, and open microsites were randomly placed at least 2 m away. The microsites were surveyed using a 0.5 m by 0.5 quadrat randomly placed beneath the shrub canopy, and the open microsites were selected within a 1.5m area of each shrub. We estimated the percent cover of residual dry matter (RDM) and of active vegetation within these quadrats. We measured the length of the longest dimension of the shrub canopy axis, its perpendicular width, and the height of each focal shrub to tip of highest green tissue. We collected all RDM within a 20 cm quadrat placed at the center of the quadrat using scissors ensuring only plants rooted within the quadrat were collected and placed it in paper bags. RDM was dried using a Yamato DKN900 drying oven at 85° C for 75 hours and then weighed to 0.001 g using Mettler Toledo analytical balance.

### Measuring ground-dwelling arthropod communities

We used pitfall traps to estimate the arthropod communities at eight pairs of shrub and open microsites at each of the nine study sites. Clear plastic drink cups (10 cm tall, 7 cm diameter) were placed in the center of a 0.5 m quadrat with the top of the cup flush with the ground. The traps were filled with a 50% propylene glycol and water mixture and were deployed for 72 hours. They were checked regularly during their deployment and were topped up with water as needed. Arthropods were sieved and placed in labelled vials containing 95% denatured ethanol. Arthropods were identified primarily to genus and family depending on the group using keys (Triplehorn and Johnson 2005, Fisher and Cover 2007, Marshall 2012) and were assigned to morphospecies within those groups. Some species are highly recognizable using certain morphological traits and known distributions such as Western black widow spiders and European earwigs and were identified to the species level. Velvet ants (Mutillidae) were not morphotyped because of strong sexual dimorphism. Only worker and major caste ants (Formicidae) were included in analyses. Springtails (Collembola) and arthropods smaller were excluded due biases arising from sieve mesh size. Larval stages and hemipteran nymphs that could not be associated with the adult form were also excluded.

### Data analysis

All statistical analyses used R version 4.0.2 (Team 2020), and the data and code are available on GitHub (https://jennabraun.github.io/rdm.gradient/).

### Biotic and abiotic drivers of arthropod community structure

We fit generalized linear mixed models (GLMM) using the R package glmmTMB (Brooks et al. 2017) to test for differences in arthropod abundance and morphospecies richness beneath the canopy of *E. californica* and open areas (n =134). One sample of 1198 individuals was excluded as an outlier in the abundance model because all of the remaining samples contained less than 350 individuals (mean = 32.3 ±39.16 SD), and 98% of the sample was comprised of a single morphospecies of *Pheidole* ant. Microsite (shrub or open), RDM biomass, RDM percent cover, deMartonne’s aridity index, and local water stress (ESI) were included as predictors. The study site was included as a random effect in both models. Only 13 quadrats contained green vegetation cover so we excluded living cover from all subsequent models. A Poisson error distribution was used to model morphospecies richness and a negative binomial error distribution was used for abundances because overdispersion was detected in the model (performance, Nakagawa et al. 2017). We compared candidate models with interactions between microsite and RDM biomass to models with only additive effects and intercept only models using AIC and likelihood ratio tests (car, Fox et al. 2012). To test if the size of the shrub influences arthropod communities, we fit a separate GLMM on the abundance and morphospecies richness for the shrub microsites samples. We used shrub width at the widest point and height as predictors and the study site as a random effect. All models were checked for co-linearity using vif (performance).

To test for differences in arthropod community composition and assess the relative importance of environment variables in structuring arthropod communities, we used canonical correspondence analysis (CCA) from the vegan package (Oksanen et al. 2010). Three a priori predictors were used: microsite (shrub or open), RDM biomass and RDM cover. The site was included to account for environmental variation. We excluded the very large outlier sample with 1198 in the CCA because this method is sensitive to extreme values (Gauch and Gauch Jr 1982). To assess the marginal significance of each hypothesized predictor, we used permutation tests with 999 permutations (anova.cca, vegan). To test for differences in species similarity between shrub and open microsites, we used analysis of similarities (ANOSIM) using Bray-Curtis distances and 999 permutations to test for significance.

### Relative interactions

To estimate the biological importance of the interactions, we calculated the effect size estimate Relative Interaction Index (RII) (Armas et al. 2004). The equation: 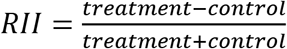 was used. Treatment is the target response value at the shrub microsite and control is the target response value at the open microsite. This measure ranges from −1 to +1, is symmetric around 0 and is common within plant ecology literature (Armas et al. 2004). Negative values indicate relative competition whilst positives indicate facilitation (Armas et al. 2004). To test for shrub facilitation for the annual communities we calculated RII using biomass and cover at the individual shrub level (n = 270 shrub/open pairs). To determine if the effects were significantly greater than zero, we bootstrapped mean and 95% confidence interval using the boot package (Canty and Ripley 2014). RII allows for different effects to be directly compared among responses on different scales and units.

Arthropod abundance and morphospecies richness RII were calculated at the site level using a bootstrap approach because several pitfall traps were disturbed in the field preventing a fully paired approach. Shrub and open microsites were paired randomly 9999 times within sites and the mean RII and SE were calculated for each iteration and site. To obtain 95% confidence intervals for the mean of all iterations we multiplied the mean standard error by 1.96. We calculated RII values for RDM biomass and RDM cover at the site level using the permutation approach to match the arthropod calculations. We then used all of the data pooled (i.e. not constrained within site) to calculate the all over RII mean and confidence intervals for both RDM and arthropod communities. To further test for a relationship between the relative effect of shrubs on the arthropod community and the annual community, we regressed each arthropod RII_abun_ and RII_richness_ and each of RDM RII_biomass_ and RII_cover_.

We tested for a relationship between environmental stress and plant productivity by regressing deMartonne’s aridity index and ESI with the mean RDM values at the site level. We then tested if the relative interactions change along the stress gradients by regressing the four RII and each of deMartonne’s aridity index and ESI at the site level.

### Morphospecies associations

To explore the regionality of significant associations between specific arthropod morphotypes and the shrub *Ephedra california,* we conducted an indicator species analysis (ISA) using the multipatt function in the R package indicspecies (De Cáceres 2013). We used regions (Panoche, Cuyama and Mojave) with microsites as the grouping factor such as Cuyama - *Ephedra*.

## Results

A total of 5868 arthropods were collected comprising 184 morphospecies in 19 orders. Ants were the most abundant group (61.6%, 3613 individuals) across 11 genera and 22 morphospecies. This was followed by bristletails (19.4 %, 1141 individuals) and beetles (6%, 346 individuals, 38 morphospecies). The full dataset is published (Braun et al, 2020).

### Biotic and abiotic drivers of arthropod community structure

Abundance and richness of arthropods were greatest beneath the canopy of the shrub *Ephedra californica* relative to open areas throughout the region (Figure 1, Table 2). RDM percent cover negatively influenced arthropod abundance without reducing richness, however, RDM biomass did not influence either arthropod community response (Table 2). The facilitation of the shrub canopy on arthropod communities was not significantly influenced by RDM cover or biomass (Appendix A3). The consistent positive effect of *E. californica* on arthropods was also not mediated by the estimates of shrub size tested (shrub width GLMMs: abundance: χ^2^ = 0.017, p = 0.895; richness: χ^2^ = 0.16, p = 0.685; shrub height GLMMs: abundance: χ^2^ = 0.568, p = 0.451; richness: χ^2^ = 1.13, p = 0.287).

**Figure 1:**
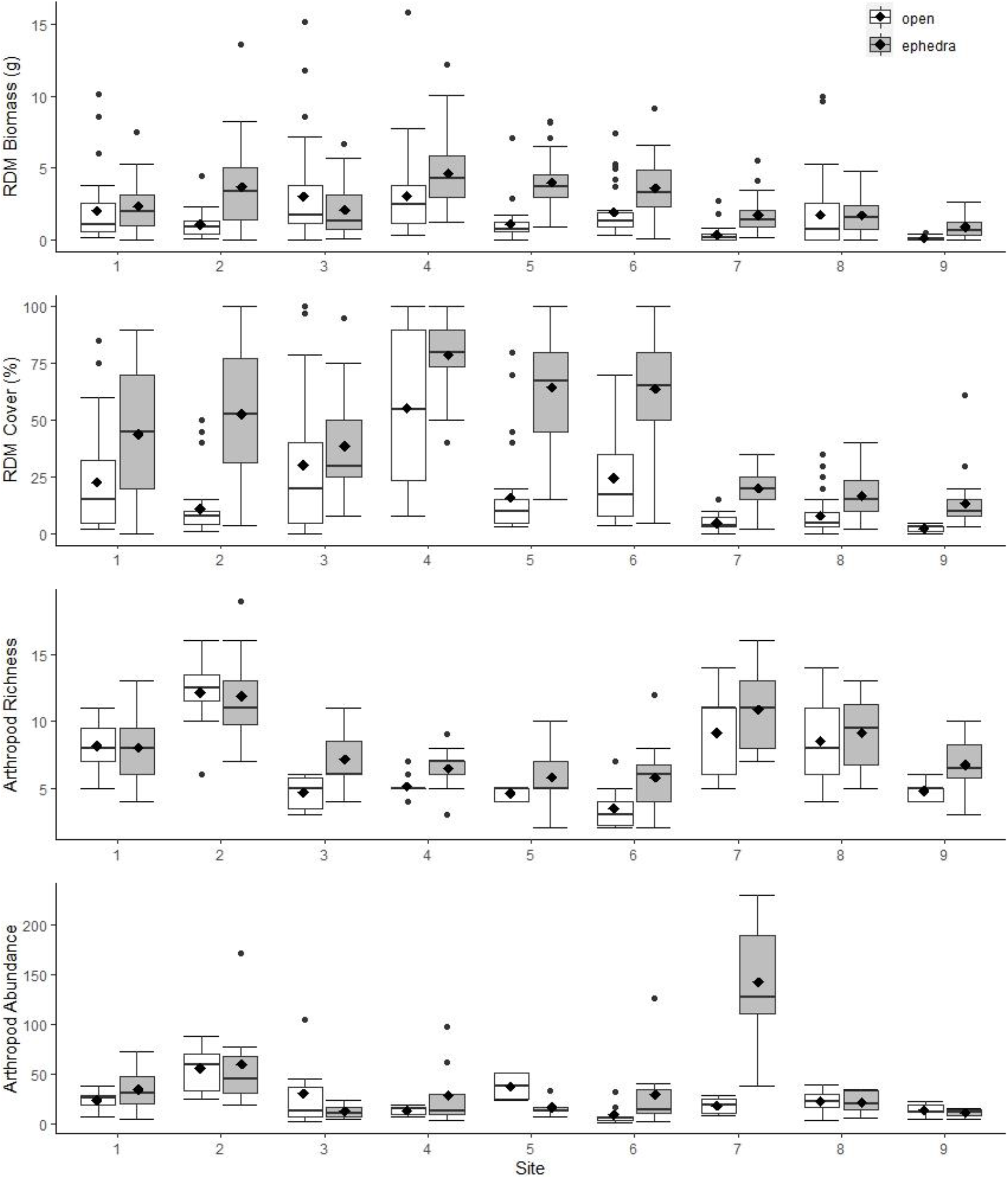
Annual plant and arthropod community responses beneath the canopy of *Ephedra californica* and in adjacent open areas. Mean responses are indicated by black diamonds; median responses are indicated by the center horizontal black line. The study sites are ordered from least arid to most arid.

**Table 2:** GLMM testing for arthropod morphospecies richness and abundance responses to microsite *(Ephedra californica* or open), residual dry matter biomass and cover, deMartonne’s aridity and local water stress (ESI). The study site was modelled as a random effect. The open microsite was used as the reference factor level, therefore, positive coefficient estimates indicate positive influence. Higher values of deMartonne’s aridity index and ESI indicate lower levels of aridity and local water stress respectively.

| | Coef | $\chi^2$ | P | | Coef | $\chi^2$ | P |
| --- | --- | --- | --- | --- | --- | --- | --- |
| Richness |  |  |  | Abundance |  |  |  |
| Shrub microsite | 0.162 | 4.846 | **0.027** |  | 0.573 | 11.652 | **0.0006** |
| RDM biomass | -0.001 | 0.005 | 0.943 |  | 0.032 | 0.913 | 0.339 |
| RDM cover | 0.0001 | 0.007 | 0.935 |  | -0.008 | 4.754 | **0.029** |
| Aridity | 0.012 | 1.286 | 0.257 |  | 0.031 | 1.872 | 0.171 |
| ESI | -2.983 | 7.601 | **0.0058** |  | -2.228 | 1.009 | 0.315 |

**Table 3:** Indicator species analysis (ISA) showing significant associations between morphotypes and *Ephedra californica.* The microsite and region were used as grouping factors to explore geographic variation in arthropod and *E. californica* associations. The five most significant associations are shown for each type.

| Morphospecies | Value | P | Panoche<br>(Semi-arid,<br>North San<br>Joaquin) |  | Cuyama<br>(Semi-arid,<br>South San<br>Joaquin) |  | Mojave<br>(Arid) |  |
| --- | --- | --- | --- | --- | --- | --- | --- | --- |
|  |  |  | Shrub | Open | Shrub | Open | Shrub | Open |
| Coleoptera: Tenebrionidae 1 | 0.389 | 0.043 | ✓ |  | ✓ |  | ✓ |  |
| Coleoptera: Curculionidae, Entiminae 1 | 0.463 |  | ✓ |  |  |  |  |  |
| Aranae: Lycosidae 1 | 0.354 | 0.022 | ✓ |  |  |  |  |  |
| Aranae: Lycosidae 3 | 0.354 | 0.025 | ✓ |  |  |  |  |  |
| Orthoptera: Rhaphidophoridae 1 | 0.612 | 0.001 |  | ✓ | ✓ |  |  |  |
| Hymenoptera: Formicidae: Solenopsis 1 | 0.655 | 0.001 | ✓ | ✓ | ✓ |  |  |  |
| Coleoptera: Tenebrionidae: Eleodes 1 | 0.601 | 0.002 | ✓ |  |  |  | ✓ | ✓ |
| Hymenoptera: Mutillidae 1 | 0.399 | 0.043 | ✓ |  |  |  | ✓ | ✓ |
| Coleoptera: Tenebrionidae 4 | 0.549 | 0.001 |  |  | ✓ |  | ✓ | ✓ |
| Hymenoptera: Formicidae: Temnothorax 1 | 0.564 |  |  |  | ✓ |  |  |  |
| Aranae: Gnaphosidae, Micaria 1 | 0.440 |  |  |  | ✓ |  |  |  |
| Aranae: Lycosidae 2 | 0.401 |  |  |  | ✓ |  |  |  |
| Aranae: Thomisidae 2 | 0.329 | 0.028 |  |  | ✓ |  |  |  |
| Aranae: Gnaphosidae 2 | 0.323 | 0.025 |  |  | ✓ |  |  |  |
| Diptera: Anthomyiidae 1 | 0.806 | 0.001 |  |  |  | ✓ |  |  |
| Hymenoptera: Formicidae: Forelius 1 | 0.682 | 0.001 |  |  |  | ✓ |  |  |
| Hymenoptera: Formicidae: Myrmecocystus 2 | 0.678 | 0.001 |  |  |  | ✓ |  |  |
| Hemiptera: Geocoridae 1 | 0.442 | 0.001 |  |  |  | ✓ |  |  |
| Orthoptera: Acrididae 1 | 0.392 | 0.005 |  |  |  | ✓ |  |  |
| Hymenoptera: Formicidae: Pogonomyrmex 1 | 0.606 | 0.001 |  |  |  | ✓ |  | ✓ |
| Coleoptera: Pyrochroidae, Cononotus 1 | 0.524 |  |  |  |  |  | ✓ |  |
| Coleoptera: Tenebrionidae 14 | 0.524 |  |  |  |  |  | ✓ |  |
| Hymenoptera: Formicidae, Crematogaster 1 | 0.511 |  |  |  |  |  | ✓ |  |
| Coleoptera: Tenebrionidae 7 | 0.468 |  |  |  |  |  | ✓ |  |
| Hymenoptera: Chyphotidae, Chyphotes 1 | 0.420 |  |  |  |  |  | ✓ |  |
| Neuroptera: Chrysopidae 1 | 0.387 | 0.008 |  |  |  |  |  | ✓ |
| Hymenoptera: Bethyridae 2 | 0.348 | 0.013 |  |  |  |  |  | ✓ |
| Lepidoptera: Micro.moth 9 | 0.341 | 0.013 |  |  |  |  |  | ✓ |
| Hemiptera: Lygaeidae 5 | 0.316 | 0.016 |  |  |  |  |  | ✓ |
| Mantodea 2 | 0.316 | 0.011 |  |  |  |  |  | ✓ |
| Microcoryphia 1 | 0.842 |  |  |  | ✓ |  | ✓ |  |

Arthropod community composition significantly varied by microsite (CCA, F = 2.54, P = 0.001), RDM cover (CCA, F = 0.178, P = 0.009) and study site (CCA, F = 2.619, P = 0.001) but not RDM biomass (CCA, F = 0.122, P = 0.169). These predictors i.e. the constrained part of the CCA explained ~22% of the measured variation in arthropod community composition. The shrub and open arthropod communities were significantly different in terms of morphospecies similarity across all sites (ANOSIM: R = 0.128, p = 0.001). Arthropod composition was also significantly dissimilar between the northern and southern semi-arid, and arid sites (ANOSIM: R = 0.5, p = 0.001).

The two estimates of environmental stress: deMartonne’s aridity index and local water stress (ESI) formed two different stress gradients during the study period. Arthropod richness increased with local water stress levels (i.e. lower ESI values) but not aridity, and arthropod abundance was not influenced by either water stress or aridity (Table 2).

### Regional morphospecies associations

The only arthropod morphospecies significantly associated with *E. californica* within all three regions was a darkling beetle in the family Tenebrionidae (Table 4). Generally, the significant associations of *E. californica* and morphospecies varied among the regions and were not dominated by any single arthropod order (Table 4). The cave cricket (Rhapidiophoridae 1) was associated with *E. californica* in the northern semi-arid sites but open areas within the southern semi-arid sites.

### Relative interactions & effects on plants

The biomass and cover of annual plant communities were consistently facilitated by *Ephedra californica* throughout the region (Figure 2, mean RII biomass = 0.30, 95% CI: 0.22 to 0.37; mean RII cover = 0.40, 95% CI: 0.34 to 0.47).

**Figure 2:**
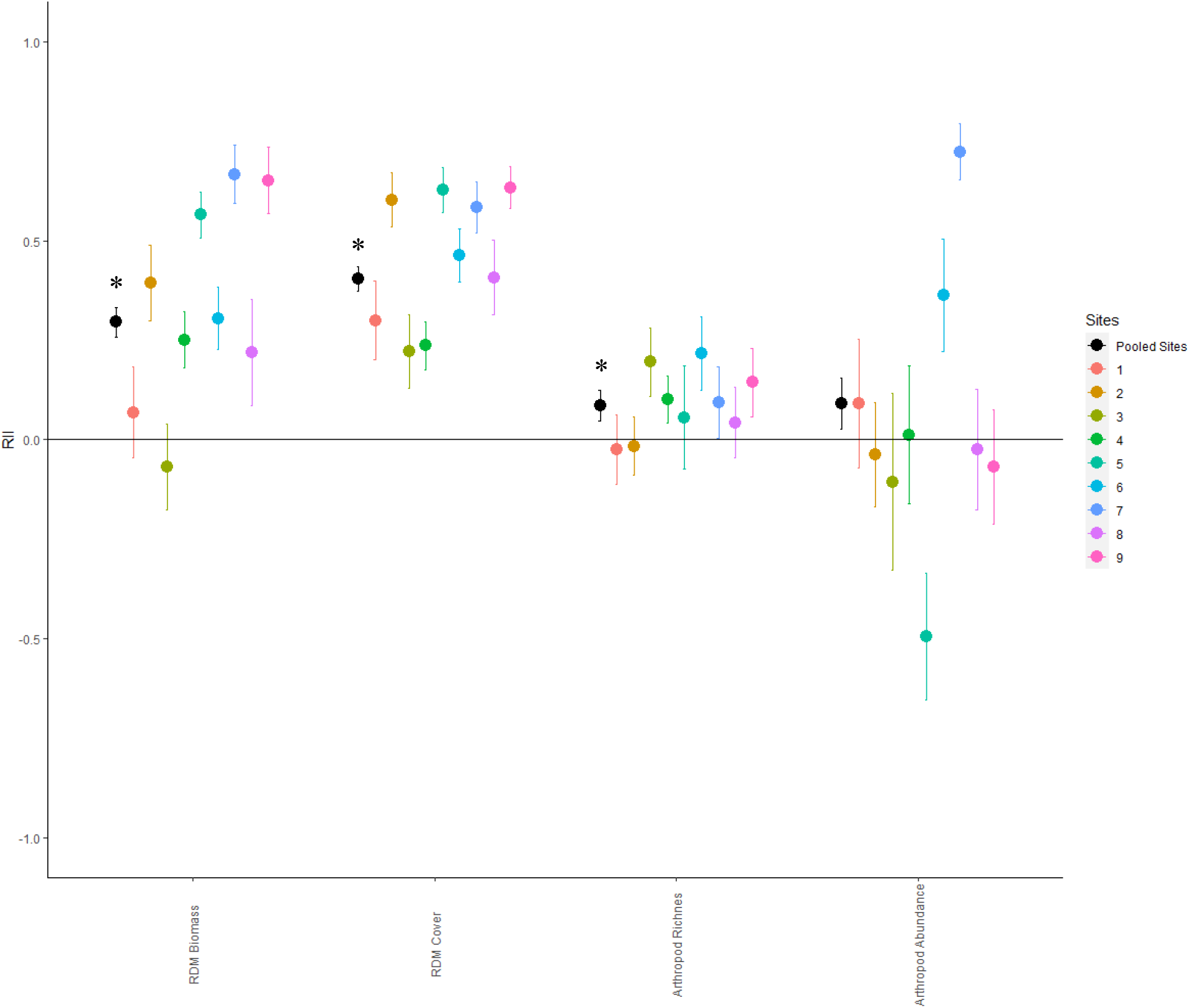
Relative interaction index (RII, bootstrapped mean ± SE) of *Ephedra californica* for each of the four community responses considered within this study. Sites are ordered from least arid to most arid. The asterisk represents a significant difference at the P < 0.05 level between RII of the pooled sites and 0.

Net effects on arthropod richness by the shrub canopy microsite were positive and significantly different from no effect (Figure 2, mean RII richness = 0.09, 95% CI: 0.01 to 0.16). Net shrub effects on arthropod abundance were not significantly different from 0 or no effect (Figure 2, mean RII abundance =0.09, 95% CI: - 0.03 to 0.22). Overall, the facilitative effects of shrubs on arthropods were independent of the facilitative effects of shrubs on annuals (Figure 3).

**Figure 3:**
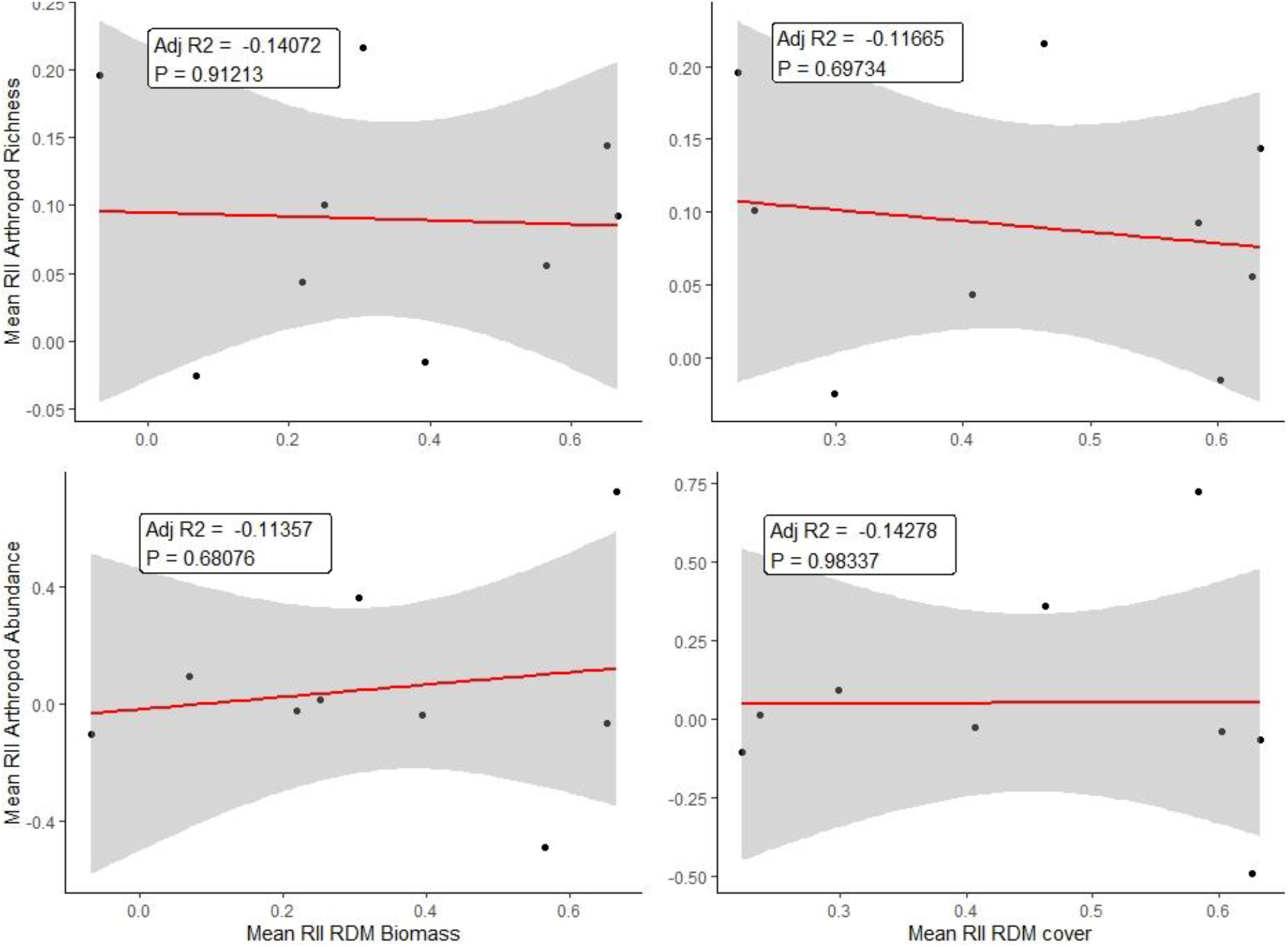
Linear regression of the mean relative interaction index (RII) for plant community response metrics (RDM biomass and cover) plotted against RII for arthropod community metrics (arthropod morphospecies richness and abundance).

Plant productivity declined along both the aridity and water-stress gradients. Arid and water-stressed sites had lower overall biomass and cover of vegetation at the soil surface as estimated by RDM (Figure 4). The relative strength of shrub facilitation did not significantly shift along either of the gradients for any of the four community response metrics measured (Figure 4).

**Figure 4:**
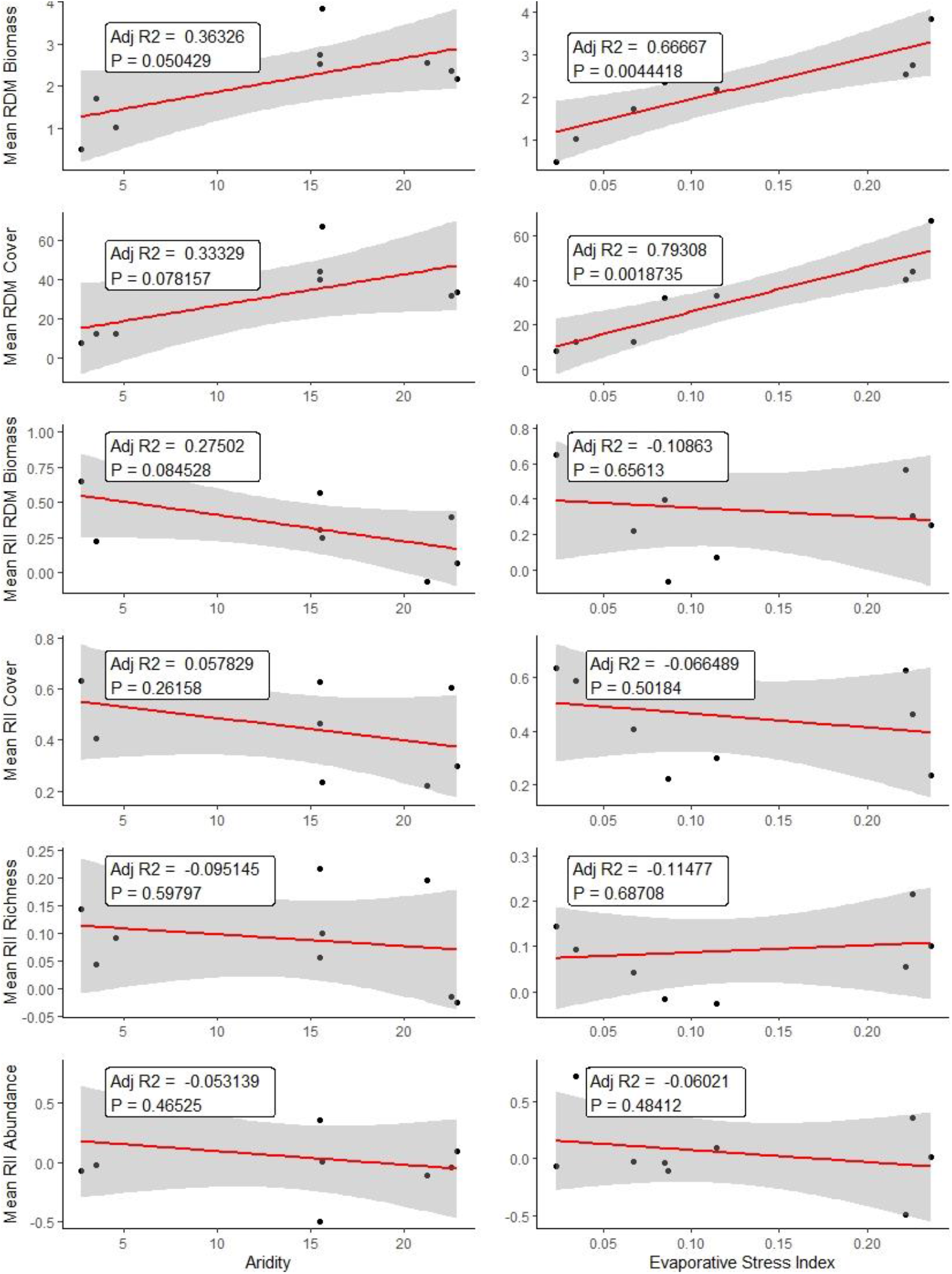
Linear regressions of deMartonne’s aridity index and evaporative stress index (ESI) plotted against mean RDM biomass and cover, and mean RII of *Ephedra californica* for each of the four community responses considered in this study. Higher values of the aridity index and ESI indicate lower aridity and water stress.

## Discussion

Plant-insect interactions are a key component of understanding biodiversity and function in many ecosystems including drylands. We used the common shrub species *Ephedra californica* and RDM to explore the relative importance of shrub canopies and ground covering vegetation i.e. RDM on the composition of resident arthropod communities. The relative effects of shrub canopies on plant and arthropod communities along an aridity gradient was also examined by exploring sites that spanned three different desert regions of Central California drylands. The hypothesis that the positive effects of *E. californica* extend to ground-active arthropods in addition to the annual plant communities was supported. Cover of RDM negatively influenced arthropod abundance and contrary to our predictions RDM biomass did not influence the strength or frequency of association by the arthropod community. The positive influence of shrubs on arthropod communities was not mediated through the understory vegetation. We propose that *Ephedra californica* is thus a foundation species regionally in Central California Deserts because it functioned positively and consistently as a benefactor for plant and arthropod communities at all environmental stress levels tested and with differences in underlying vegetation.

Shrubs likely provide a generalized facilitation mechanism such as microclimate amelioration; however, biotic interactions in addition to amelioration are important to structuring arthropod communities. Within Cuyama and Mojave, bristletails (Microcoryphia) were significantly associated with *E. californica*. Bristletails are important detritivores and feed on decaying organic matter (Triplehorn and Johnson 2005). This suggests indirect effects of the shrub mediated through the annual layer but on a different time scale than measured here. Trophic interactions within arthropod food webs are likely important to the composition at fine spatial scales. Wolf spiders (Lycosidae) were strong indicators of *E. californica* at the semi-arid sites. Wolf spiders are important predators and are active hunters that stalk prey. In grasslands, wolf spiders are major consumers (20%) of secondary biomass (Van Hook Jr 1971). Spiders are generally considered ecological indicators (Maelfait et al. 1989, Hendrickx et al. 1998). *Chyphotes* (Hymenoptera: Chyphotidae), were previous placed within Bradynobaenidae (Pilgrim et al. 2008) and were also associated with Mojave Desert *E. californica*. This group has also been previously placed within the family Mutillidae and very little is known about their biology. Previous work in the Mojave desert that focused on the foundation species creosotebush (*Larrea tridentata)* reported that Chyphotidae were exclusively associated with shrubs (Ruttan et al. 2016). Thus, these finding show that both of these foundation species are important to this group which again suggests generalized positive effects of desert shrubs on arthropods. The variation in significant associations shows that *E. californica* supports a wide range of arthropod functional groups from predators to detritivores that then provide a suite of ecosystem services locally (Whitford 2000, Prather et al. 2013). This species of shrubs and other desert species such as *Larrea tridentata* are foundational because they very broadly support biodiversity – i.e. diversity in very different levels of taxa.

Variation on gradients at regional scales are important to examine to better understand species pools and also the relative importance of drivers at those scales such as climate. At the community level, decreases in primary productivity can be a surrogate to estimate environmental stress (Grime 1977, Armas et al. 2011). The plant community RDM biomass and cover declined with both the aridity and local water stress gradient. Aridity and water-stress did not influence the abundance of arthropods, and richness was higher at the more water-stressed sites. Therefore, we did not detect the expected effects of stress on the arthropod communities. The stressors measured here may not be directly relevant to the arthropod communities at these scales because they are relatively motile. Additionally, aridity and water stress were estimated at relatively larger scales and vegetation responded but at fine-scales the arthropods may be sampling and experiencing climate at fine scales instead (Sylvain et al. 2014). The frequency of shrub facilitation on either the plant or arthropod communities was always positive and did not increase with increasing stress. The discrepancy between our results and other tests of the SGH may be due to the net seasonal estimates of RDM used here. The RDM was the pooled annual community productivity for the full growing season and therefore it may be necessary to refine by plant life stage to test SGH on that temporal scale. Similar to the findings here, most papers that do not support SGH still tend not shift to competition with stress in many ecosystems (He et al. 2013). Shrubs and arthropods occupy different trophic levels and ecological niches, and are therefore unlikely to be in direct competition for space, light, or resources (Schoener 1974). These findings highlight the importance of shrubs to arthropod communities because facilitation was consistent throughout the gradient and at least estimated by the net final productivity did not indirectly depend on other vegetation present in the system.

Habitat for all species is all ecosystems including deserts is a global issue. Maintaining heterogeneity in rangelands can lead to more positive outcomes including supporting biodiversity and enhancing ecosystem functioning (Fuhlendorf and Engle 2001). Shrubs create heterogeneity by creating variation in physical structure and the evidence here suggests that shrubs can be an important form of habitat for the annual plant and arthropod communities of drylands. However, these traits that make *E. californica* beneficial to ecosystems through heterogeneity may also support the facilitation of invasive species. The invasive *Bromus* grass forms thick monocultures and preferentially establishes underneath *E. californica* shrubs (Lucero et al. 2019). Our results suggest that the net balance of shrub effects on arthropod abundance is neutral due the strong facilitation of ground cover. Bare ground is necessary nesting habitat for many insects including solitary native bees (Michener 2000) and larval antlions (Myrmeleontidae) require unvegetated ground to build pits to hunt (Triplehorn and Johnson 2005). RDM can be used a measure of interference on arthropod communities, but we do not know mechanisms until tested experimentally. RDM also interferes with the movements of the endangered blunt-nosed leopard lizard (*Gambelia sila)* (Filazzola et al. 2017). Therefore, a potential consequence of the intensifying invasion by non-native ground-covering plant species is that the positive effects of shrubs on arthropods and vertebrates can be reduced.

In drylands, shrubs are foundation species that stabilize the environment for other taxa including plant and arthropod communities. Ground-active arthropods provide a suite of ecosystem services such nutrient cycling, seed dispersal and pest control (Whitford 2000). Maintaining *Ephedra californica* cover through healthy canopies and not necessarily the largest individual sized shrubs will support both ecosystem function and biodiversity in dryland ecosystems (Lortie et al. 2018). The focus of facilitation research on plants and vertebrates means that the importance of these foundation species to dryland ecosystems has likely been underestimated.

## Data Statement

Arthropod data openly available at KNB (doi:10.5063/F1J101JB), all code and data available at https://github.com/jennabraun/rdm.gradient.

## Appendix

**A1:**
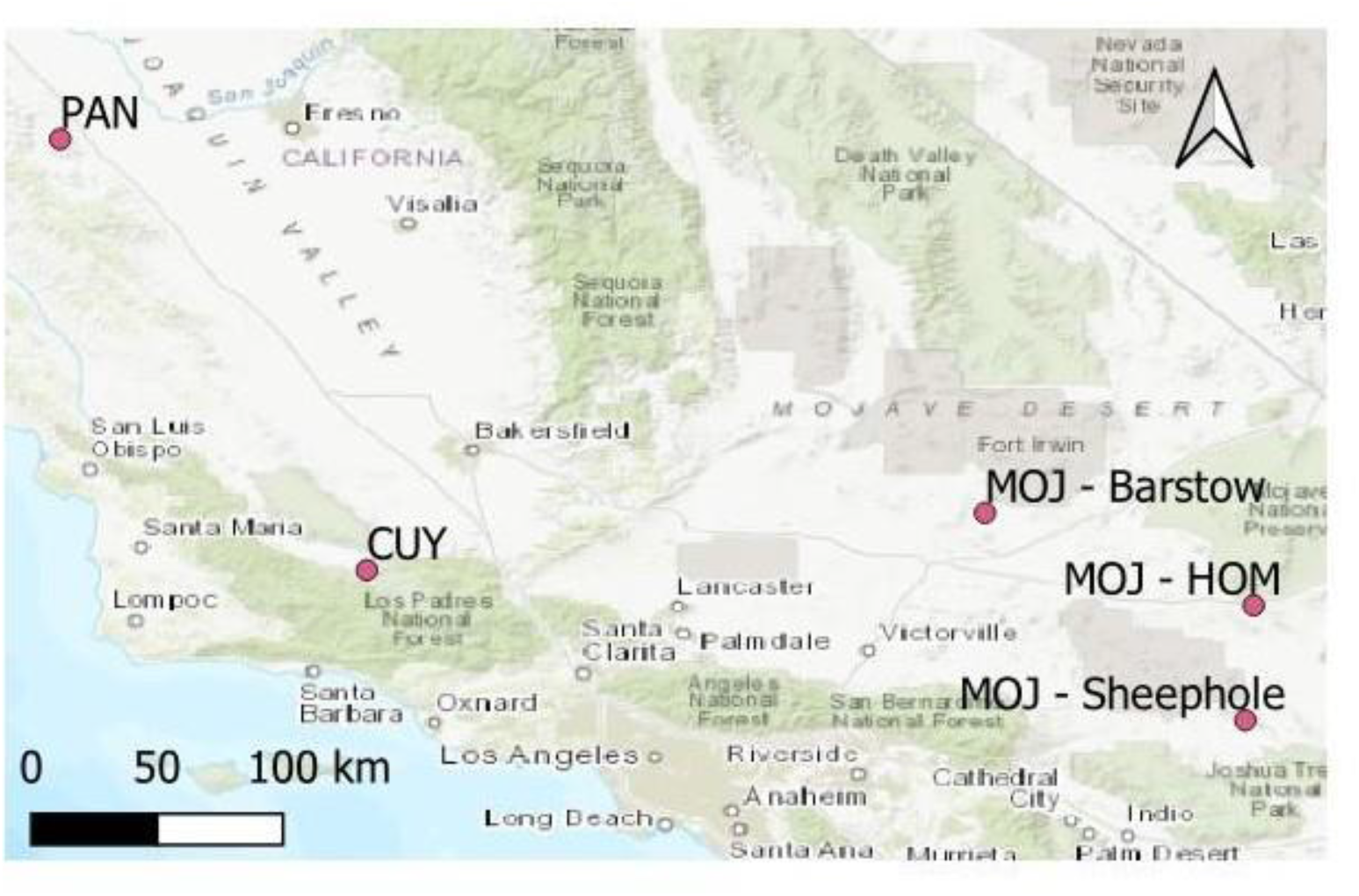
The study sites were located in the San Joaquin Desert of California, USA: Panoche Hills (sites 4-6) and Cuyama Valley (sites 1-3), and the Mojave Desert: Heart of the Mojave, Sheephole Valley and near Ft. Irwin (sites 7-9).

**A2:** Dates, times and orbit identification data for EcoSTRESS sensor corresponding to data used in this project.

| ID | Latitude | Longitude | Date and time | Orbit Number | Build ID | Scene ID |
| --- | --- | --- | --- | --- | --- | --- |
| 1 – C1 | 34.84873 | -119.483 | 2019-06-30 18:51:58 UTC | 5579 | 601 | 9 |
| 2 – C2 | 34.85362 | -119.486 | 2019-06-30 18:51:58 UTC | 5579 | 601 | 9 |
| 3 – C3 | 34.93824 | -119.481 | 2019-06-30 18:51:58 UTC | 5579 | 601 | 9 |
| 4 – PAN3 | 36.70000998 | -120.801 | 2019-06-21 14:56:35 UTC | 5437 | 601 | 1 |
| 5 – PAN2 | 36.70554 | -120.812 | 2019-06-21 14:56:35 UTC | 5437 | 601 | 1 |
| 6 – PAN1 | 36.70654 | -120.812 | 2019-06-21 14:56:35 UTC | 5437 | 601 | 1 |
| 7 - M2 | 34.698199 | -115.684 | 2019-07-01 18:02:52 UTC | 5594 | 601 | 11 |
| 8 – M1 | 34.20568 | -115.72 | 2019-07-01 18:02:52 UTC | 5594 | 601 | 11 |
| 9 – M3 | 35.094051 | -116.835 | 2019-07-01 18:02:52 UTC | 5594 | 601 | 11 |

A3: AIC scores comparing models with interaction terms between RDM biomass and microsite (shrub or open), and RDM cover and microsite (shrub or open) to additive models. The interaction terms were insignificant in all models. Chisquare tests were used (anova) to test if the interactive models were significantly different from the additive model.

**A3:**
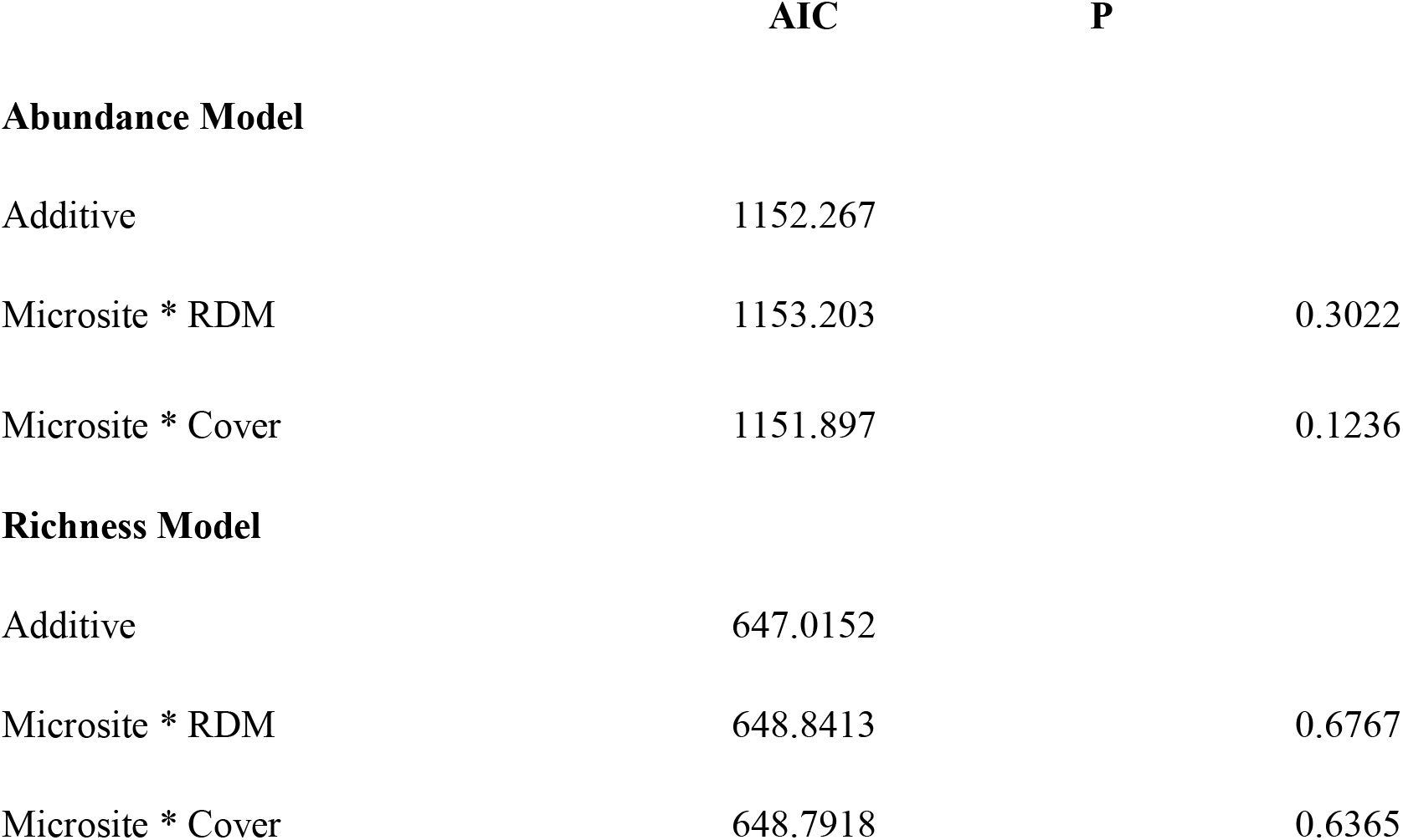
AIC scores comparing models with interaction terms between RDM biomass and microsite (shrub or open), and RDM cover and microsite (shrub or open) to additive models. The interaction terms were insignificant in all models. Chisquare tests were used (anova) to test if the interactive models were significantly different from the additive model.

|  | AIC | P |
| --- | --- | --- |
| **Abundance Model** |  |  |
| Additive | 1152.267 |  |
| Microsite * RDM | 1153.203 | 0.3022 |
| Microsite * Cover | 1151.897 | 0.1236 |
| **Richness Model** |  |  |
| Additive | 647.0152 |  |
| Microsite * RDM | 648.8413 | 0.6767 |
| Microsite * Cover | 648.7918 | 0.6365 |

